# Growth and extended survival of *Escherichia coli* O157:H7 in soil organic matter

**DOI:** 10.1101/235275

**Authors:** Gitanjali NandaKafle, Amy A. Christie, Sébastien Vilain, Volker S. Brözel

## Abstract

Enterohaemorrhagic *Escherichia coli* such as serotype O157:H7 are a leading cause of food-associated outbreaks. While the primary reservoir is associated with cattle, plant foods have been associated as sources of human infection. *E. coli* is able to grow in the tissue of food plants such as spinach. While fecal contamination is the primary suspect, soil has been underestimated as a potential reservoir. Persistence of bacterial populations in open systems is the product of growth, death, predation, and competition. Here we report that *E. coli* O157:H7 can grow using the soluble compounds in soil, and characterize the effect of soil growth in the stationary phase proteome. *E. coli* 933D (stxII^-^) was cultured in Soil Extracted Soluble Organic Matter (SESOM) and the culturable count determined for 24 d. The proteomes of exponential and stationary phase populations were characterized by 2D gel electrophoresis and protein spots were identified by MALDI-TOF mass spectrometry. While LB controls displayed a death phase, SESOM grown population remained culturable for 24 d, indicating an altered physiological state with superior longevity. This was not due to decreased cell density on entry to stationary phase as 24h SESOM populations concentrated 10-fold retained their longevity. Principal component analysis showed that stationary phase proteomes from SESOM and LB were different. Differences included proteins involved in stress response, motility, membrane and wall composition, nutrient uptake, translation and protein turnover, and anabolic and catabolic pathways, indicating an altered physiological state of soil-grown cells entering stationary phase. The results suggest that *E. coli* may be a soil commensal that in absence of predation and competition maintains stable populations in soil.

## Introduction

*Escherichia coli* O157:H7 and related enterohaemorrhagic strains have been associated with many serious food-associated outbreaks, (Hilborn et al. 1999; Currie et al. 2007; Grant et al. 2008; King et al. 2009). The infectious dose is low so that food products are required to be free from enterohaemorrhagic *Escherichia coli* O157:H7, but despite various measures taken during processing, consumers can still be exposed to this pathogen (LeBlanc 2003; Yang et al. 2017). Cattle are widely believed to be the primary host and several outbreaks have been associated with beef-based products (Currie et al. 2007; King et al. 2009). *E. coli* O157:H7 is known to be associated with the bovine gastrointestinal tract, specifically the cecum (Yoon and Hovde 2008; Wang et al. 2017), currently believed to be the primary source of entry into the food chain. More recently various plant foods such as spinach, tomato, lettuce and fresh fruits have been identified as sources (Grant et al. 2008; Herman et al. 2015; Denis et al. 2016). Initially these foods were thought to be fecally contaminated, but recent reports suggest growth of *E. coli* O157:H7 (Brandl 2008; Wright et al. 2013; Wright et al. 2017) in tissues of salad leaves and tomatoes. Upon inoculation from an unknown source, the enteric bacteria multiply inside the growing plant, and cannot be removed through surface treatment such as washing. The annual nature of these crop plants excludes them as an environmental reservoir of these enteric bacteria. Rather, these crop plants would need to be infected during growth.

*E. coli* are found in both gastrointestinal systems, and in the environment (Adamowicz et al. 1991; Ishii et al. 2006; Ksoll et al. 2007). Once shed from a mammalian host, *E. coli* populations are widely believed to enter a dead end, relying for extended survival on stress responses (Winfield and Groisman 2003). The paradigm assumes slow decline following fecal contamination, the basis of the fecal coliform test (Tallon et al. 2005). This is supported by decline of *E. coli* O157:H7 in manure (Williams et al. 2008; Looper et al. 2009) and in soil (Berry and Miller 2005) over time. Yet enterohaemorrhagic *E. coli* maintain culturable populations in various soils for many months, even when moisture limited, and with slower decline at lower temperatures (Berry and Miller 2005; Fremaux et al. 2008). Some *E. coli* appear to grow in sub-tropical environments such as riverbank soil and river sediment (Desmarais et al. 2002). More recently persistent *E. coli* populations have been reported from temperate forest, watershed soils (Byappanahalli et al. 2006; Ishii et al. 2006), and pasture (NandaKafle et al. 2017). Naturalized *E. coli* strains believed to be autochthonous to soil were able to maintain populations in soils (Jang et al. 2017).

Persistence of bacterial populations in soil would require a suitable nutrient pool. Soil is a complex assemblage of particulate components with varying concentrations of organic and inorganic matter. The dissolved organic matter (DOM) in soils is a cocktail of sugars, aromatic compounds, amino acids, and organic and fatty acids between C_14_ and C_54_ (Huang et al. 1998; Kalbitz et al. 2000). The concentrations of solutes like amino acids range from 0.1-5 μM. Monoprotic acids (e.g. formate, acetate and lactate) range from 1 μM to 1000 μM, and di- and trivalent low molecular organic acids (e.g. oxalate, malate and citrate) from 0.1-50 μM (Strobel 2001; Pizzeghello et al. 2006). Monomeric intermediates such as carboxylic acids and amino acids have residence times in the order of hours in soils (Jones et al. 2005; Van Hees et al. 2005). Carbohydrates like mono-, di- and oligosaccharides vary in presence and concentration (Lynch 1982; Guggenberger and Zech 1993b; a; Kaiser et al. 2001; Kalbitz et al. 2003). Surprisingly, glucose is present in soils up to 100 μM concentrations (Schneckenberger et al. 2008). The variety of sugars, organic and amino acids in these soils suggest that enteric bacteria should, generally, be able to grow here. We have reported a detailed analysis of liquid extract of deciduous forest soil, able to support growth of *Salmonella* Typhimurium (Liebeke et al. 2009).

The source of contamination of annual food crops by enterrohaemorrhagic *E. coli* is unresolved. Soil has been underestimated as a potential reservoir. As persistence of bacterial populations in open systems is the product of growth, death, predation, and competition, measurement of numbers over time shows the overall net effect, and cannot inform autecology of the species. Whether population maintenance of *E. coli* O157:H7 in soils is due to a combination of cell division and death, predation and competition, or simply to extended survival alone, is unresolved. It has been shown that *E. coli* O157:H7 is able to grow in sterile fresh water (Vital et al. 2008). In order to understand how soil is a potential reservoir for enterrohaemorrhagic *E. coli*, autecological studies are required. Here we report that *E. coli* O157:H7 is able to grow in liquid extract of soil. Furthermore, soil extract-grown populations demonstrate extended culturability over cultures grown in laboratory media, and display a unique stationary phase proteome.

## Materials and Methods

### Culture and culture media

The partially attenuated *E. coli* O157:H7 933D (*stx*-II) (Strockbine et al. 1986) was maintained in 50% (v/v) glycerol at −80°C. Soils used were corn field soil (Brandt silty clay loam, Aurora, South Dakota, USA), a commercially available garden top soil, and deciduous forest soil (Oak Lake Field Station, Brookings, South Dakota, USA). Cow manure from a herd fed an antibiotic-free diet was obtained from the South Dakota State University Beef Unit. Soil-extracted solubilized organic matter (SESOM) was prepared as described previously (Vilain et al. 2006). Briefly, 100g of air-dried soil, or 90g soil and 10g manure, was suspended in 500 mL MOPS buffer (10 mM, pH 7, 50°C) and kept shaking at 200 rpm for 1h. The extracts were filtered sequentially through filter paper, hydrophilic PVDF membranes with 5, 1.2, and 0.45, pore sizes to remove particulates, and sterilized using a polyethersulfone membrane with a 0.22 μm pore size. The sterility of each batch was determined by placing 5 μl SESOM onto LB agar plates and incubating at 30°C for 24h.

### Culturing conditions

Growth and survival in the various liquid extracts was determined by measuring the optical density periodically. Overnight cultures of *E. coli* O157:H7 were prepared in LB broth, diluted 1:1,000 into 50mL fresh LB broth and incubated to mid-exponential phase at 28°C (3h, A_546_ = 0.41). Cells were harvested by centrifugation (10,000 × g, 10 min, 30 °C), washed twice, and re-suspended in 2mL sterile tap water. Triplicate 250mL flasks with 50mL pre-warmed liquid medium (LB, 1/40^th^ strength LB and SESOM from deciduous forest soil) were inoculated to an initial A_546_ of 0.005, and incubated at 28°C while shaking (120 rpm). The culturable count was determined every hour till 8h, at 24h and daily till 24d by the droplet plate technique (Lindsay and Von Holy 1999). Briefly, 20 μL volumes of serial dilutions were plated onto LB agar and incubated for 18h at 30°C.

### Effect of cell density on culturability

The effect of cell density on extended culturability of populations was investigated by concentrating or diluting populations, and re-suspending in cell-free supernatants of the same culture type. LB-grown populations were harvested at 24h of incubation (10,000 × g, 10 min, 30 °C) and re-suspended to one tenth their density in cell-free supernatant. Conversely, 1/40^th^ strength LB and SESOM-grown populations were re-suspended to ten-fold density in their respective cell-free supernatants. All cultures were then incubated at 28 °C while shaking, and the culturable count determined every 24h to day 24.

### Protein sample preparation

SESOM and LB – grown populations of *E. coli* O157:H7 were harvested in mid-exponential phase (180min and 140min) at A_546_ 0.05 5 and 0.183, respectively, and SESOM, LB, and 1/40^th^ strength LB populations were harvested in late stationary phase (3d). Cells were harvested by centrifugation (10,000 × g, 10 min, 4°C), washed in 5mL potassium phosphate buffer (100 mM, pH 7.0), and re-suspended in 2ml IEF buffer (7 M urea, 2 M thiourea, 2% (w/v) 3-[3-chloamidopropyl] dimethylammonio-1-propanesulfonate (CHAPS), 2% (w/v) Amidosulfobetaine-14 (ASB14), 10mM dithiothreitol (DTT) and 2% (v/v) carrier ampholytes (pH 3.5 – 10; Amersham)). Cells were disrupted by two cycles of freeze thaw (from −80°C to 20°C) followed by ultrasonication at 4°C (15W, 12 pulses of 3min). Cell debris was removed by centrifugation (10,000 × g, 10min), and the protein concentration was determined using the Bradford protein assay (BioRAD), with bovine serum albumin as the standard (Vilain et al. 2001).

### Two-dimensional gel electrophoresis (2DE)

IPG strips (pH 4 – 7, 18 cm, GE Healthcare) were re-hydrated for 16h with 400 μL IEF buffer containing 50 μg protein for 2D gel map construction, and 200 μg protein per IPG strip for protein identification. Proteins were separated by IEF on an Amersham Pharmacia horizontal electrophoresis system for a total of 44 kVh (150 V for 1 h, 350 V for 1 h, 500 V for 4 h, 750 V for 1h, 1 kV for 1 h, 1.5 kV for 1 h and 3.5 kV for 11 h). After IEF, the IPG gel strips were frozen at −80 °C, thawed and equilibrated for 10 min in equilibration buffer (6 M urea and 30% glycerol, 1% SDS) with 20mg/mL DTT, and for 10 min in equilibration buffer with 260 mM iodoacetamide. The second dimension consisted of SDS-PAGE using a 12.5% (w/v) running polyacrylamide gel and a 4.65% stacking gel (width, 18 cm; length, 20 cm; thickness, 1 mm). Gels were stained with silver (Rabilloud 1992) for spot detection and protein map construction, and with colloidal Coomassie Blue G250 for protein identification (Vilain et al. 2001). Uninoculated SESOM was run on a one-dimensional SDS PAGE to check for proteins present, but following staining, none were found.

### Gel analysis, spot detection and protein map construction

Gels were scanned using a transmission scanner (ScanMaker 9800XL, Microtek) in transmission mode. Gel images were analyzed using PDQuest software (version 7.3.1; Bio-Rad) which allows detection, quantification and matching of protein spots. Spots were quantified on a Gaussian image and pooled on a reference image. The following formula was used to calculate the quantity of Gaussian spot: Spot height × σ_x_ × σ_y_ × π; where: Spot height is the peak of the Gaussian representation of the spot, σ_x_ is the standard deviation of the Gaussian distribution of the spot in the direction of the x axis, and σ_y_ is the standard deviation in the direction of the y axis. SESOM-derived spots either higher than two-fold or less than half the intensity in LB broth were excised from stationary phase LB, LB 1/40, and SESOM derived gels, and identified by MALDI-TOF mass spectrometry of tryptic digests as described previously (Voigt et al. 2006), but using the *E. coli* O157:H7 EDL933 sequence database (ftp://www.expasy.org/databases/complete_proteomes/fasta).

Principal component analysis (PCA) of the 2D electrophoretograms was performed as described previously (Vilain and Brozel 2006), using Statgraphics Plus 4.0 (Manugistics). Briefly, calculations of the Eigen value were comprised by taking the data set and subtracting the mean value from each dimension (ie. effect of culture medium) until all means were zero. A covariance matrix was then calculated since the data set has more than one dimension. By calculating the covariance matrix on means of zero a line develops that characterizes the data. The lines, or Eigen values, determine the statistical significance of each of the components.

## Results

The enterohaemorrhagic pathogen *E. coli* O157:H7 was able to grow using water-soluble organic matter from various soils, as indicated by increases in optical density during incubation (Fig. 1). The yield in SESOM was 1 Log lower than in LB broth, and varied among extracts of various soils. Nutrient carry-over from the LB-pre-culture was avoided by extensive washing and inoculating to a low initial density of A_546_ = 0.005. These results indicate that, in the absence of competition and predation, populations of *E. coli* O157:H7 933D should be able to grow and divide in soils, as supported by population increases in the various SESOM evaluated. Populations in LB broth started loosing culturability after 3 d of incubation, with long-term stationary phase of 1% of the population density setting in on d9, as is well established. Surprisingly, SESOM-grown populations remained culturable for 24 d, the duration of the experiment (Fig. 2a). The observed loss of culturability in LB was not due to an adverse pH, measured as 6.8 (d16) and 7.3 (d19). The pH of uninoculated SESOM was 6.8, decreasing to 5.9 (d16) and 5.8 (d19).

**Fig. 1.**
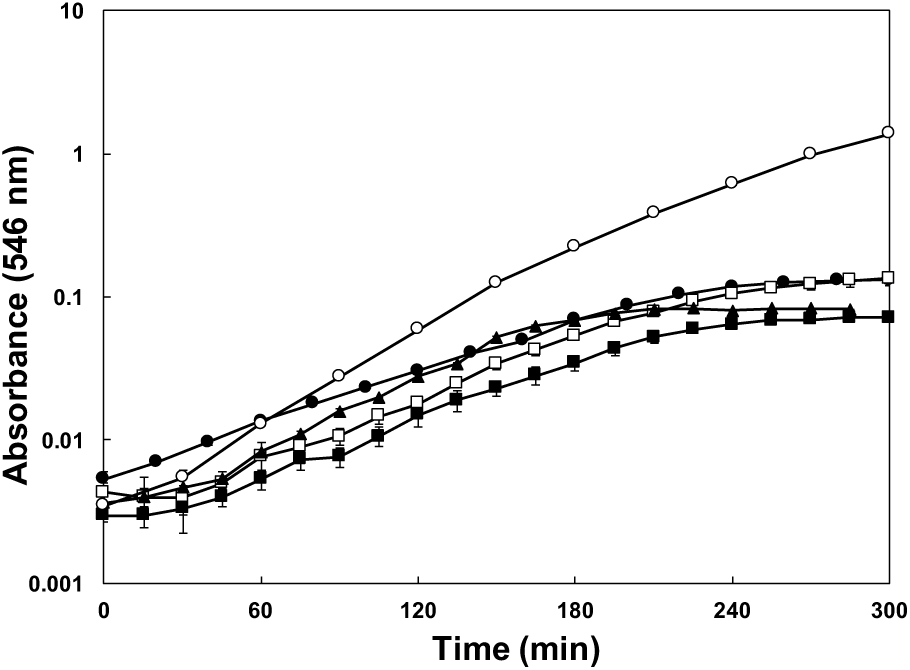
Growth of *E. coli* 0157:H7 933D stxII-in SESOM from deciduous forest soil (●), corn field soil (■), corn field soil supplemented with 10% (m/v) cow manure (□), garden soil (▲), and LB broth (○) while shaking at 30°C.

**Fig. 2.**
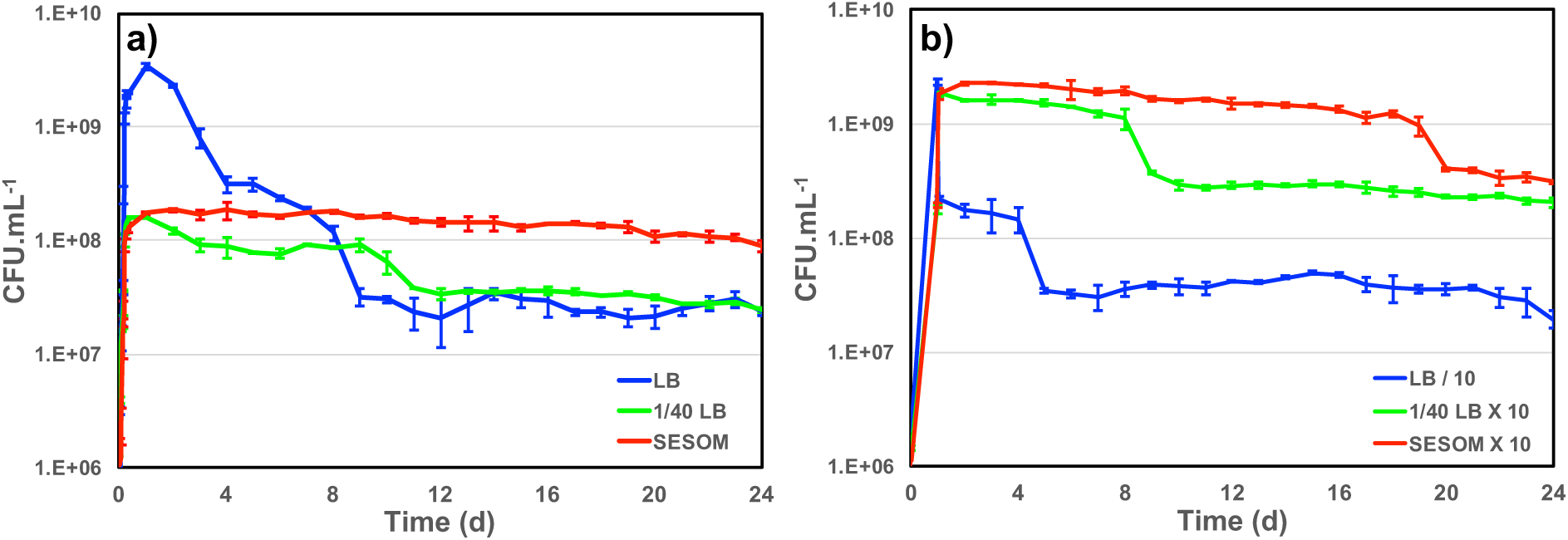
Growth and survival of *E. coli* O157:H7 933D in LB, dilute LB (1/40) and SESOM (a), and when cultures were either concentrated ten-fold in own supernatant (SESOM and 1/40th LB – grown), or diluted ten-fold (LB – grown) at 24h (b). Error bars indicate one standard error of the mean.

To determine whether cell density during entry into stationary phase affects future culturability, we sought to culture in LB to the same population density achieved in SESOM. Various dilutions of LB (1/30, 1/40, 1/50, 1/70, and 1/100) were evaluated to determine which supported a final optical density similar to SESOM (results not shown). LB diluted 40 times yielded the desired density, and the resulting populations remained culturable longer than in LB, with a stable population of 10^8^ CFU/mL at d9 (Fig. 2a), after which culturability declined. This indicated that population density in stationary phase may play a role in maintenance of culturability of cells, with higher density associated with decreased survival. The pH on d12 was 6.6. Cells that entered stationary phase due to nutrient limitation in 1/40^th^ strength LB were more resilient than populations grown to higher density in LB. SESOM grown cells were, however, more resilient than 1/40^th^ LB grown cells, although both entered stationary phase due to nutrient limitation (Fig. 2a). This indicated that soil grown *E. coli* populations would persist longer in soil than predicted by laboratory experiments. Cultures in M9 minimal medium with 10 g.L^−1^ glucose displayed loss in culturability over time, similar to in LB (data not shown), indicating that increased longevity could not be attributed to growth requiring a greater degree of anabolic reactions.

Cell density appeared to play a role in stationary phase survival of LB-grown populations (Fig. 2a). To further investigate the role of cell density in survival, we modified cell density 10-fold upon entry into stationary phase. LB-grown stationary phase cells (24h) were harvested and resuspended to one tenth their original density in their own spent broth, and the culturable count determined for 24d. The population lost one log_10_ of culturability after d4 (Fig. 2b), as opposed to 2 log_10_ in undiluted culture (Fig. 2a). This could indicate that LB-grown *E. coli* are able to maintain only a certain cell density into stationary phase. To determine whether the resilience of populations grown in 1/40 strength LB was due to lower final density, stationary phase populations were concentrated ten-fold and resuspended in their own supernatant. The increased cell density did not initially lead to much loss of culturability, similar to the un-concentrated culture (Fig. 2 a & b). After decline at d8, late stationary phase population density remained at ten-fold that of the original 1/40 th LB grown culture. Importantly, the concentrated 1/40^th^ LB population was at the same density as LB-grown population entering stationary phase, but did not undergo the 2 log_10_ decline. Thus 1/40^th^ LB – grown cells were more likely to survive than LB-grown ones, irrespective of cell density post-stationary phase. These results indicated that conditions upon entry into stationary phase affect the condition of the cells, thereby determining their potential for survival over long term incubation.

SESOM-grown populations maintained at ten-fold concentration in their own spent medium declined slowly, only showing a five-fold decline at d19 (Fig. 2b). Thus SESOM – grown cells were more likely to survive than 1/40^th^ and LB-grown ones, irrespective of cell density post-stationary phase. Collectively the results indicated that extended longevity of SESOM-grown populations was due to both a lower cell density, but also to a SESOM-associated factor. These results suggested that SESOM-grown populations had an altered physiological state when entering stationary phase.

To gain insight into possible physiological reasons underlying the extended longevity of SESOM-grown populations, the proteomes of LB and SESOM-grown cultures in exponential and stationary phase (d3), were determined by 2DE, and compared to the 1/40^th^ LB stationary phase proteome. For exponential phase, care was taken that populations had not yet begun transition to stationary phase. The five proteomic datasets were then subjected to principle component analysis (PCA). Four components were revealed at an Eigen value greater than 1, *viz*. 2.69, 2.51, 1.53 and 1.20. These components were sequentially compared in pair-wise fashion using biplots (Fig. 3). The results showed that exponential phase LB- and SESOM grown populations differed significantly, as did stationary phase populations in the two media. Intriguingly, the LB 1/40^th^ stationary proteome was very similar to the SESOM-proteome in the first three of four coordinates, and quite different to the LB stationary phase proteome. Collectively the PCA analysis showed that stationary phase populations had culture medium-specific proteomes that could explain the different physiological states and propensity to survive.

**Fig. 3.**
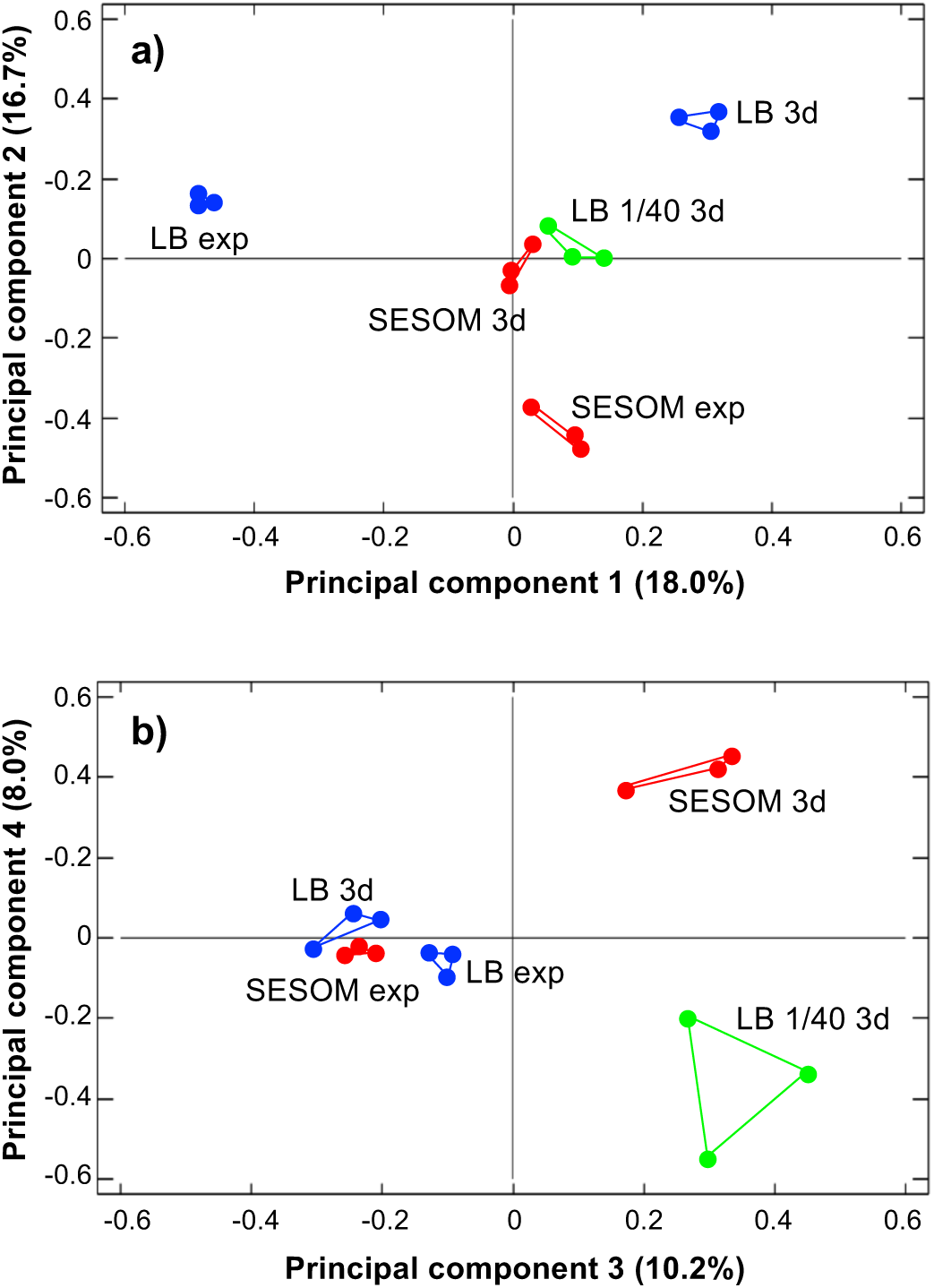
Principle component analysis of exponential and stationary phase (3d) proteomes of *E. coli* O157:H7 933D cultured in LB, 1/40strength LB and SESOM at 30°C.

A large number of protein spots had significantly different abundance as determined using the criteria outlined above in materials and methods. All these spots were identified by MALDI-TOF MS, and collectively paint a unique physiological state of *E. coli* O157:H7 persisting in soil organic matter (Table 1). Stationary phase LB populations appeared to experience several stresses as indicated by elevated levels of the universal stress protein UspA and the carbon starvation protein Slp. They also had elevated levels of the alkyl hydroperoxide reductase AhpC. By contrast SESOM-grown cells appeared less stressed and more active, indicated by increased levels in transcriptional (DksA and RpoA) and translational proteins (GroEL, TufA and YeiP). This suggested sustained transcriptional and translational activity during stationary phase in SESOM versus LB. Many uptake systems were either over or under-expressed in SESOM-grown populations, including outer membrane and periplamic uptake systems. This indicates that cells growing on SESOM have the ability to sense what the surrounding environment has to offer. These proteins reinforce the notion that *E. coli* O157:H7 is very adaptable to nonhost environments such as soil. In addition to various uptake systems, several systems involving substrate metabolism were found to be over and under-expressed in SESOM-grown cells, indicating different approaches to catabolic activity in stationary phase. Structurally, cells grown in SESOM appear to be different based on the expression of several membrane and cell structure proteins, primarily those involved in membrane lipid biosynthesis. YmcD and Adk were both up-regulated in SESOM. This suggests that the cellular envelope is formed differently in SESOM-grown populations as opposed to LB-or LB 1/40-grown populations. Perhaps the cellular envelope is thicker to provide protection from adverse conditions. Both structural and regulatory flagellar proteins were present in increased abundance in SESOM-grown cells suggesting that the cells are potentially motile and responsive to chemotactic behavior in soil organic matter. Overall, SESOM-grown stationary phase cells appeared less stressed, more motile, metabolically different, and with suggestions of less altered membrane composition when compared to LB-grown populations.

**Table 1.**
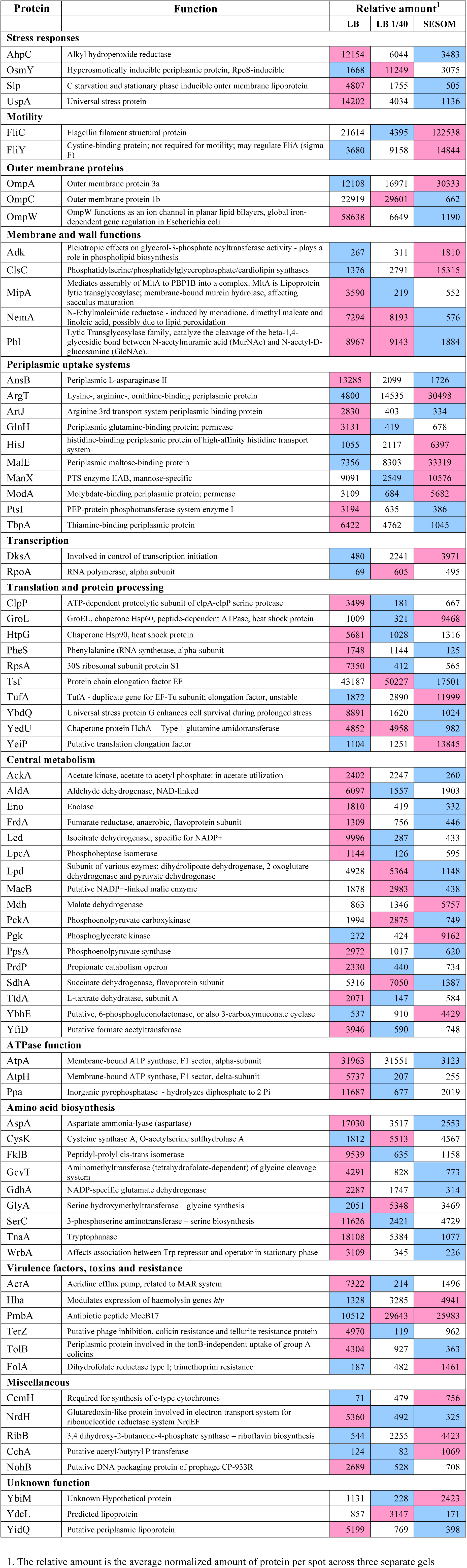
Proteins of different abundance in stationary phase (3d) populations of *E. coli* O157:H7 grown and maintained in LB, 1/40-strength LB and SESOM at 30 °C.

## Discussion

*E. coli* is not thought to survive for long periods outside the host intestine, so produce-associated outbreaks have widely been ascribed to recent fecal contamination. The suspected sources of produce contamination include soil amendments (manure or compost), irrigation water contaminated with cattle feces, or contaminated surface runoff (Ongeng et al. 2015). Our results showed that *E. coli* O157 can grow using nutrients available in soils (Fig. 1). There have been countless studies reporting numbers of *E. coli* O157 in soils over time, and some have suggested growth in soil. Survival of *E. coli* in soil has been reported by many researchers; more than 200 d under natural environmental conditions and 500 d in frozen soil and on plant roots (Gagliardi and Karns 2002; Islam et al. 2004). This is the first report showing definitively that *E. coli* O157 is able to grow using water soluble nutrients in soil.

Soil-grown *E. coli* O157 appeared more resilient than laboratory-grown cultures, with almost 100% culturability maintained over 28 d (Fig. 2). This finding pointed to an altered physiological state of SESOM-grown cells entering stationary phase. This suggests that *E. coli* responds differently to nutrient limitation in SESOM, preparing cells for stationary phase differently. To gain insight into possible physiological reasons underlying the extended longevity of SESOM-grown populations, the proteomes of LB and SESOM-grown stationary phase cultures were compared (Table 1). The stationary phase proteome of SESOM-grown *E. coli* differed significantly from LB-grown and dilute LB-grown populations (Fig. 3), indicating cells with substantially altered composition, and therefore catalytic and structural properties.

SESOM grown cells had lower levels of several proteins associated with cellular responses to stress, including Alkyl hydroperoxide reductase (AhpC), the carbon starvation response lipoprotein Slp, and the universal stress protein UspA (Table 1).AhpC is the primary degrader of hydrogen peroxide and reactive nitrogen intermediates in *E. coli,* protecting the cell against oxidative stresses (Chen et al. 1998). The substantially lower concentration of AhpC in SESOM-grown cells indicated decreased oxidative stress, or possible alternative mechanisms to cope with reactive oxygen and nitrogen species. Slp accumulates in response to carbon starvation (Alexander and St John 1994), but our data showed that LB grown cells expressed the most Slp, although SESOM-grown cells were clearly nutrient starved following entry in stationary phase (Liebeke et al. 2009). Markedly, SESOM-grown cells responded differently to nutrient starvation and entry in stationary phase. UspA is induced as soon as the growth rate falls below the maximum rate supported by the medium (Nystrom and Neidhardt 1994). Despite the abrupt transition from exponential to stationary phase in SESOM, cells expressed less UspA than in LB. SESOM populations contained a much greater amount of the flagellar components FliC and FliY, indicating increased motility.

OmpA, the major outer membrane protein in *E. coli*, was more prevalent in the SESOM population. Loss of OmpC in *E. coli* contributes to antibiotic resistance (Liu et al. 2012), but this is only significant in the exponential-phase, while such difference in stress-resistance becomes trivial after bacteria reach the stationary phase (Wang 2002). The elevated OmpA in LB populations is likely due to the high NaCl concentration. The ratio of OmpC to OmpF increases at higher temperature and pH, as well as under oxidative stress (Snyder et al. 2013), consistent with increased level of AhpC in LB cells. Elevated levels of Adk and ClsC indicated differences in membrane lipid composition of SESOM versus LB-grown populations due to their role in synthesis of phospholipids. An addition, Adk has been linked to mutational fitness effects. It was also observed that the length of the lag phase is more sensitive to variation in Adk catalytic capacity than is the exponential growth rate, so that the lag phase appears to be optimal with respect to variation in Adk catalytic capacity (Adkar et al. 2017). NemA, abundant in LB cultures, is involved in reductive degradation of toxic nitrous compounds (Umezawa et al. 2008), again consistent with elevated AhpC in LB populations. Enhanced levels of the peptidoglycan-modulating factors MipA and Pbl in LB cultures indicates differences in cell wall structure between the stationary phase cultures. High levels of MipA have been reported in sessile compares to planktonic cultures (Rivas et al. 2008).

Periplasmic nutrient uptake systems varied in quantity across the three culture media, but would be remnants from exponential phase where amino acid and sugar uptake were required. The high levels of AnsB, ArtJ and GlnH in LB cultures is puzzling as LB supplies ample amino acids derived from tryptone and yeast extract. Our forest SESOM did not contain detectable levels of lysine, arginine, ornithine or histidine (Liebeke et al. 2009), explaining the enhanced level of ArgT and HisJ. ArgT expression is increased in response to nitrogen starvation and during early response glucose limitation (Kabir et al. 2004; Franchini and Egli 2006), and our SESOM contained very little glucose. Molybdenum (molybdate) is essential as cofactor for the assembly and function of several enzymes including nitrate reductase, formate dehydrogenase, dimethyl-sulfoxide reductase, trimethylamine-N-oxide reductase, and biotin-sulfoxide reductase (Rajagopalan and Johnson 1992). PtsI, more prevalent in LB, is a component of the glucose uptake system that is inhibited by α-ketoglutarate during nitrogen limitation, when it was over expressed the metabolic rate was increased fourfold (Chubukov et al. 2017).

DksA was highly expressed in SESOM grown cells, suggesting a stringent response with induction of ppGpp synthesis due to nutrient limitation. DksA activated by ppGpp binds to the β-subunit of RNA polymerase, directly affecting the affinity to different promoters and thus altering the expression level of more than 80 genes, most importantly suppression of all components of the protein biosynthesis system: rRNA, ribosomal proteins, and translation factors (Pletnev et al. 2015). This indicates that transcription is shut down tightly in SESOM-grown stationary phase cells.

LB populations appeared more stressed as indicated by elevated levels of various stress proteins and chaperones. LB populations contained more ClpP, part of the proteosomal protein degradation system. Controlled degradation of cytoplasmic proteins has long been considered essential for survival of bacteria under stress conditions, due to the requirement for efficient removal of misfolded or otherwise damaged proteins by ClpP (Weichart et al. 2003). The corresponding low abundance of ClpP in cultures with no decline phase indicated either a reduced need for protein turnover, or a lower degree of damaged proteins. A different profile of damaged proteins was supported by the differences in chaperones GroL (SESOM and 1/40 LB) and HtpG (LB). HtpG expression is increased in cells grown in a complex medium with ample amino acid availability (LB) following heat shock, but low in glucose minimal medium. HtpG expression unaffected or even repressed by imposition of a nutrient stress condition in minimal medium (Mason et al. 1999). The stressed nature of LB stationary cells was supported by elevated levels of Tsf (Elongation factor EF), which plays a role in sequestering surfaces of heterologous proteins to prevent protein-protein interactions leading to formation of inclusion bodies (Han et al. 2007). Elevated levels of the putative stress proteins YbdQ and YedU further indicated greater degree of stress in LB populations, as also indicated by alkyl hyodroperoxide reductase. The elevated levels of TufA (Elongation factor Tu) in SESOM indicates minor starvation due to nutrient limitation. TufA plays an important role in a minor starvation defense mechanism where it helps in rescuing stuck ribosomes (Pletnev et al. 2015).

A total of 17 central carbon metabolism enzymes were detected, and 14 of these were more abundant in LB, indicating that SESOM populations had prepared for reduced metabolic activity going into stationary phase. *E. coli* grown in rich medium undergo a reconstruction of their proteome in stationary phase, with increases in proteins required for scavenging and metabolizing rare nutrients and general cell protection (Li et al. 2014). An example was AckA (acetate kinase), part of the acetate switch that occurs as cells deplete their environment of acetate-producing carbon sources and scavenge for acetate. The accumulation of extracellular acetate during stationary phase occurs as cells co-metabolize acetogenic amino acids, e.g. l-threonine and l-alanine, with those that require the TCA cycle, e.g., L-glutamate (Wolfe 2005). A second example was the elevated levels of ATP synthase components in LB populations, indicating continued need for ATP synthesis driven by periplasmic proton motive force. A third example was PckA (phosphoenolpyruvate carboxykinase), elevated in LB and 1/40^th^ LB. PckA increases 100-fold in the stationary phase independent of cyclic AMP, probably to provide carbohydrates required for energy reserves after cessation of growth, since protease activity, Krebs cycle enzyme activities, and glycogen synthesis all increase in the stationary phase (Goldie and Sanwal 1980).

LB-grown populations had higher overall levels of amino acid biosynthetic enzymes, than both SESOM and dilute LB cultures. This contrasts with the abundance of metabolizable oligopeptides available in LB. However, bioassay of LB medium after growth of *E. coli* showed that it no longer contains significant amounts of recoverable L-serine, L-threonine, L-proline, glycine, L-arginine, L-glutamine, L-asparagine, L-cysteine, and L-lysine (Sezonov et al. 2007), indicating a need for synthesis in stationary phase.

*E. coli* O157 933D appears well adapted to grow using soluble nutrients available in soil (SESOM). Moreover, SESOM grown populations did not display a detectable death phase, but remained culturable for at least 24 d. This was supported by the substantially altered proteome of SESOM-grown stationary phase populations. Our results suggest that *E. coli* may well be a soil commensal that maintains stable populations in soil, as growth supported by soil nutrients combined with enhanced longevity of cells would help counter the effects of competition and predation. Soil itself should, therefore, be included as potential source of contamination of fresh produce. Future work should investigate the roles of competition and predation affecting *E. coli* populations in soil.

## Acknowledgements

We thank David Francis for donating *E. coli* O157:H7 933D, and Birgit Voigt of the Institute for Microbiology, Ernst Moritz Arndt University, Greifswald, Germany for protein identification. GN and AC were supported by the South Dakota Agricultural Experiment Station. This research was supported by the South Dakota Agricultural Experiment Station. We acknowledge use of the SDSU-FGCF supported in part by NSF/EPSCoR Grant No. 0091948 and by the State of South Dakota.

